# Neuronal activity induces aggrecan expression to drive perineuronal net formation in cortical neurons

**DOI:** 10.1101/2025.05.29.656756

**Authors:** Kentaro Nakayama, Ayumu Mubuchi, Shinji Miyata

## Abstract

Synaptic plasticity, driven by activity-dependent changes in neuronal connectivity, underlies learning and memory. However, the mechanisms that sustain long-term synaptic stabilization are not well understood. Perineuronal nets (PNNs), which are lattice-like extracellular matrix structures that enwrap neurons, are thought to stabilize synapses. However, the mechanisms regulating their formation remain unclear. Here, we found that neuronal activity induces transcription of aggrecan, a core PNN component, which in turn promotes PNN assembly. In primary cultured mouse cortical neurons, pharmacological stimulation of neuronal activity robustly increased aggrecan expression, whereas activity blockade suppressed it. Among the PNN-related genes examined, aggrecan alone exhibited strong activity-dependent transcriptional regulation. Analysis of published *in vivo* datasets revealed selective upregulation of aggrecan in parvalbumin-positive interneurons following sensory stimulation. This regulation required calcium influx through voltage-gated calcium channels and was dependent on cAMP response element binding protein (CREB) signaling and chromatin remodeling. These findings reveal a novel molecular link between neuronal activity and PNN formation, providing insights into the mechanisms underlying the long-term stabilization of neural circuits.

## Introduction

Neural circuits reorganization in response to external stimuli is a critical process for learning and memory. How transient activation of neurons leads to long-term changes in neural circuit properties is a fundamental question in neuroscience. Neuronal activity-dependent gene expression allows neurons to translate transient patterns of activity into long-term changes in connectivity and function. Over the past three decades, research has shown that activity-dependent gene expression supports synaptic plasticity, dendritic remodeling, and the establishment of an inhibitory/excitatory balance, all of which are essential for learning, memory, and adaptive behavior ^1–7^.

Activity-dependent gene expression can be divided into two temporal categories: immediate-early genes (IEGs) and late-response genes (LRGs). IEGs are rapidly activated within minutes of neuronal stimulation and regulate subsequent transcriptional programs. Prominent examples include *Fos* and *Npas4*, which encode transcription factors that drive the expression of LRGs ^8–10^. LRGs include regulators of structural and functional synaptic plasticity. This temporal difference highlights the complexity of activity-dependent transcriptional regulation, with IEGs acting as molecular first responders to activity changes and LRGs involved in long-term neural circuit remodeling.

Previous studies have established the importance of calcium signaling pathways in activity-dependent gene expression. Membrane depolarization following synaptic activation opens L-type voltage-gated calcium channels (L-VGCCs), resulting in a rapid influx of calcium ions into the neuron. Calcium also enters through N-methyl-D-aspartate (NMDA) receptors. Calcium signaling activates downstream molecules, including calcium/calmodulin-dependent protein kinases (CaMKs), protein kinase A (PKA), and calcineurin, which modulate transcription factors through phosphorylation or dephosphorylation ^3,7^. For example, phosphorylation of cAMP response element binding protein (CREB) drives the transcription of IEGs such as *Fos* and *Npas4*. Moreover, neuronal activity maintains long-term gene expression patterns required for synaptic plasticity and memory formation through epigenetic mechanisms. Specifically, neuronal activation leads to dynamic and genome-wide changes in DNA methylation and chromatin accessibility, as well as enhancer activation via histone acetylation^8,9,11–14^.

Although activity-dependent gene expression has been extensively studied in the context of synaptic plasticity and circuit remodeling, how such gene expression affects the extracellular matrix (ECM) surrounding neurons remains poorly understood. Perineuronal nets (PNNs) are lattice-like ECM structures that preferentially surround parvalbumin-positive inhibitory neurons (PV neurons) in the cerebral cortex ^15–18^. Hyaluronan (HA), a long linear glycosaminoglycan, serves as the scaffold for PNNs ^19^. A single HA filament binds several HA-associated chondroitin sulfate proteoglycans (CSPGs), such as aggrecan, versican, neurocan, and brevican, to form a macromolecular complex in the extracellular space. The interaction between CSPGs and HA is stabilized by HA and proteoglycan-link proteins (HAPLNs) ^20,21^. The core HA-CSPG complex is further cross-linked by tenascin-R and protein tyrosine phosphatase receptor type Z1 (PTPRZ1) ^22,23^.

Many studies have shown that PNN formation drives the transition from the highly plastic juvenile neural circuits to the more stable circuits characteristic of the adult brain. For example, critical period plasticity, typically restricted to the juvenile brain, can be reactivated in the adult brain by enzymatic digestion of PNNs ^24^. In addition, removal of PNNs impairs both memory acquisition and storage, highlighting their essential role in the stability of neural circuits ^25–30^. Since the PNNs are crucial for determining the stability of neural circuits, it is possible that PNNs, like synapses, are not static structures, but rather undergo dynamic remodeling in response to external stimuli. Indeed, sensory deprivation has been shown to reduce PNN density in somatosensory and visual cortices ^24,31,32^. In addition, excessive depolarization enhanced PNN formation, whereas blockade of depolarization suppressed PNN formation both *in vitro* and *in vivo* ^33–35^. Neuronal activity has been shown to promote the expression of some PNN components, such as aggrecan, brevican, and HAPLN1 ^32,36–38^. However, the molecular mechanisms that regulate the gene expression of PNN components in an activity-dependent manner are largely unknown.

Using primary cultured mouse cortical neurons, this study comprehensively analyzed transcriptional changes of PNN components upon pharmacological modulation of neuronal activity. Among several PNN components, only aggrecan, encoded by the *Acan* gene, showed robust expression changes in response to neuronal activity. We also found that activity-dependent expression of *Acan* requires calcium influx via L-VGCC and activation of CREB. Additionally, our results suggest that chromatin remodeling of the *Acan* promoter region is essential for activity-dependent PNN formation.

## Results

### Neuronal activity promotes *Acan* expression and PNN formation in cortical neurons

To investigate whether neuronal activity regulates PNN formation and the expression of PNN-related genes, we isolated primary cortical neurons from embryonic day 17 (E17) mouse brains and maintained them *in vitro* for 13-15 days (DIV13-15) (Fig. 1a). Depolarization was induced by application of 25 mM potassium chloride (KCl) for 24 hours. We then assessed PNN formation by staining for *Wisteria floribunda* agglutinin (WFA), a marker of PNNs. Quantification of WFA intensity around PV neurons revealed a significant increase in the KCl-treated group compared to controls (Fig. 1b, c), indicating that increased neuronal activity promotes PNN formation.

**Figure 1.**
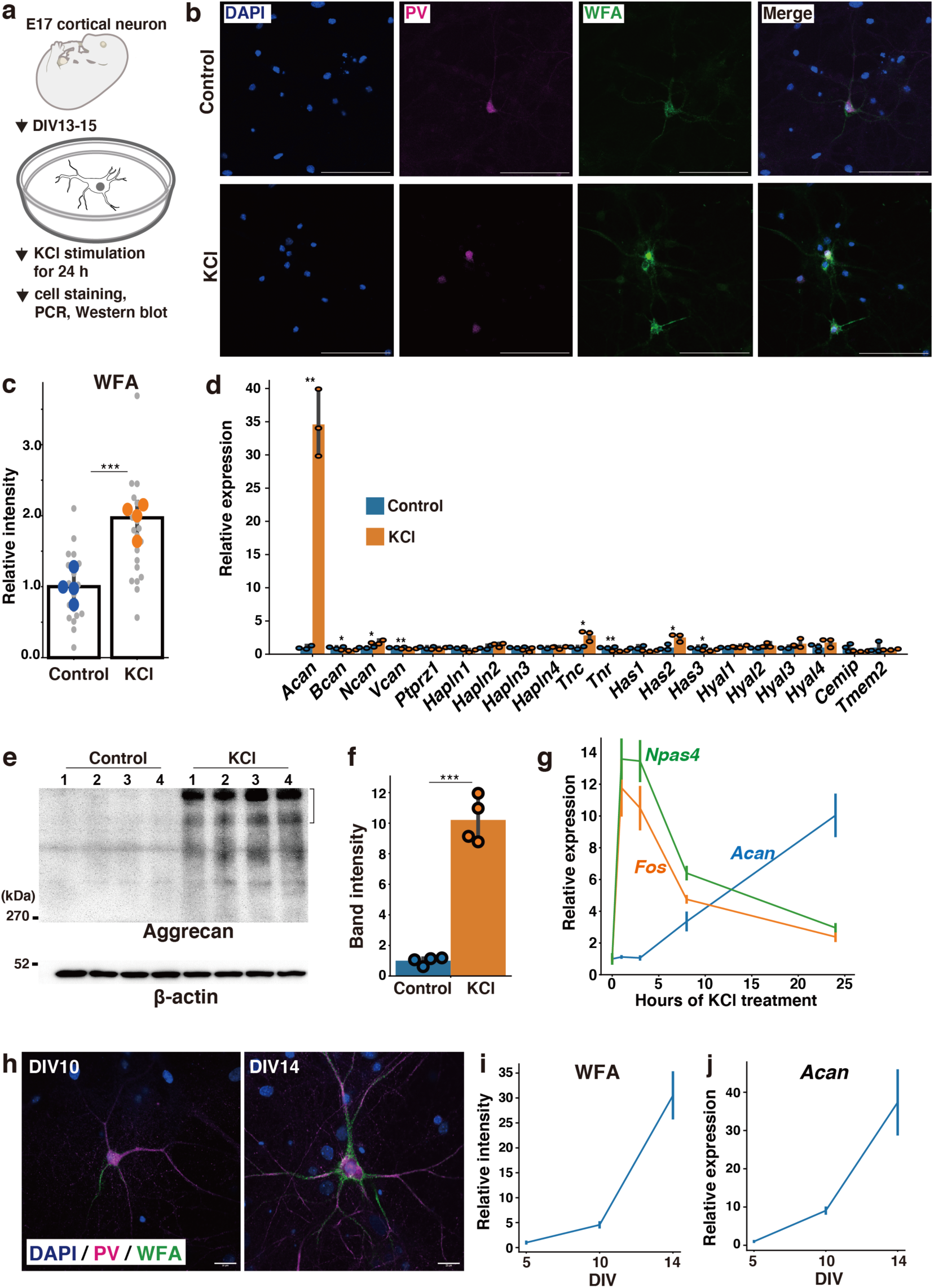
Neuronal activity promotes *Acan* expression and PNN formation in cortical neurons. (**a**) Schematic representation of the experimental design. (**b**) Representative images showing WFA-positive PNNs around PV neurons in control and KCl-treated (25 mM, 24 h) cultures at DIV14. (**c**) Relative WFA intensity around PV neurons. Gray dots represent individual images from each replicate, and blue (control) and orange (KCl) dots indicate the mean WFA intensity per well (n = 4 wells). (**d**) Expression levels of PNN-related genes in control and KCl-treated cultures at DIV16, as determined by quantitative PCR. Values are expressed relative to control (set to 1) (n = 3 wells). (**e**) Western blot analysis of aggrecan and β-actin in control and KCl-treated cultures. (**f**) Band intensity of aggrecan shown in (e), normalized to β-actin and expressed relative to control (n = 4 wells). (**g**) Expression changes of *Acan*, *Fos*, and *Npas4* at the indicated time points after KCl treatment (n = 4 wells). (**h**) Representative images showing the developmental formation of WFA-positive PNNs around PV neurons at DIV10 and DIV14. (**i**) Relative WFA intensity at DIV5, DIV10, and DIV14. Values are normalized to DIV5 (n = 4 wells). (**j**) Expression levels of *Acan* at DIV5, DIV10, and DIV14. Values are normalized to DIV5 (n = 4 wells). All statistical tests are two-tailed Student’s t-tests. Data are presented as mean ± SEM. ***P<0.001, **P<0.01, *P<0.05. Scale bars are 100 μm.

To identify the transcriptional changes associated with KCl-induced PNN formation, we analyzed the expression of PNN-related genes by quantitative PCR. The genes examined included CSPGs (*Acan*, *Bcan*, *Ncan*, *Vcan*, *Ptprz1*), link proteins (*Hapln1-4*), tenascins (*Tnc*, *Tnr*), hyaluronan synthases (*Has1-3*), and hyaluronidase-related genes (*Hyal1-4*, *Cemip*, *Tmem2*). Among these, *Acan* showed a dramatic increase upon KCl treatment (Fig. 1d). None of the other PNN-related genes showed robust changes, indicating that *Acan* is selectively upregulated in response to neuronal activity. Consistent with the transcriptional changes in *Acan*, Western blot analysis confirmed that aggrecan was significantly increased by depolarization at the protein level (Fig. 1e, f).

To gain insight into the cellular mechanisms underlying activity-dependent *Acan* expression, we compared the temporal transcriptional dynamics of *Acan* with those of IEGs such as *Fos* and *Npas4*. *Fos* and *Npas4* showed a rapid and transient increase 1 h after depolarization, whereas *Acan* expression was induced late and showed significant upregulation only after 8 h, which lasted for 24 h (Fig. 1g). These results suggest that *Acan* is not an IEG, but is induced as part of a delayed transcriptional program. We next examined whether the temporal dynamics of *Acan* expression coincide with the timing of PNN formation during neuronal maturation *in vitro*. WFA-positive PNNs were scarcely observed at DIV5. However, from DIV10 to DIV14, WFA staining intensity progressively increased and PNNs became clearly detectable around PV neurons (Fig. 1h, i). Notably, the expression pattern of *Acan* paralleled PNN formation: compared to DIV5, *Acan* mRNA levels increased approximately eightfold at DIV10 and thirty-fivefold at DIV14 (Fig. 1j).

### Activity-dependent *Acan* expression in PV neurons in the mouse visual cortex

To determine whether neuronal activity upregulates the expression of PNN-related genes *in vivo*, and to identify the neuronal populations expressing PNN-related genes, we analyzed a published transcriptomic dataset ^39^. This dataset contains the gene expression profiles of four neuronal populations in the visual cortex: EMX1-positive excitatory neurons and PV-, VIP-, and SST-positive inhibitory neurons. These populations were collected before and after light exposure, which stimulated neuronal activity (Fig. 2a). PNN-related genes showed distinct expression patterns in the four neuronal subtypes. (Fig. 2b, Supplementary data 1). *Acan*, *Hapln1*, *Hapln4*, and *Cemip* were selectively expressed in PV neurons, which are enveloped by PNNs (Fig. 2b). Furthermore, *Acan* expression was increased approximately 2-fold in PV neurons after light exposure, consistent with our *in vitro* findings. While IEGs such as *Fos* and *Npas4* are rapidly and transiently upregulated within 1 hour after light exposure (Supplementary data 1), *Acan* expression showed a delayed induction, again consistent with *in vitro* results. Among other CSPGs, *Bcan* and *Ncan* were highly expressed in EMX1-positive excitatory neurons, while *Vcan* and *Ptprz1* were enriched in VIP-positive inhibitory neurons; however, their expression showed only modest increases in response to visual stimulation compared to *Acan* (Fig. 2b). Taken together, these results suggest that increased neuronal activity leads *Acan* expression specifically in PV neurons, thereby promoting PNN formation around these cells.

**Figure 2.**
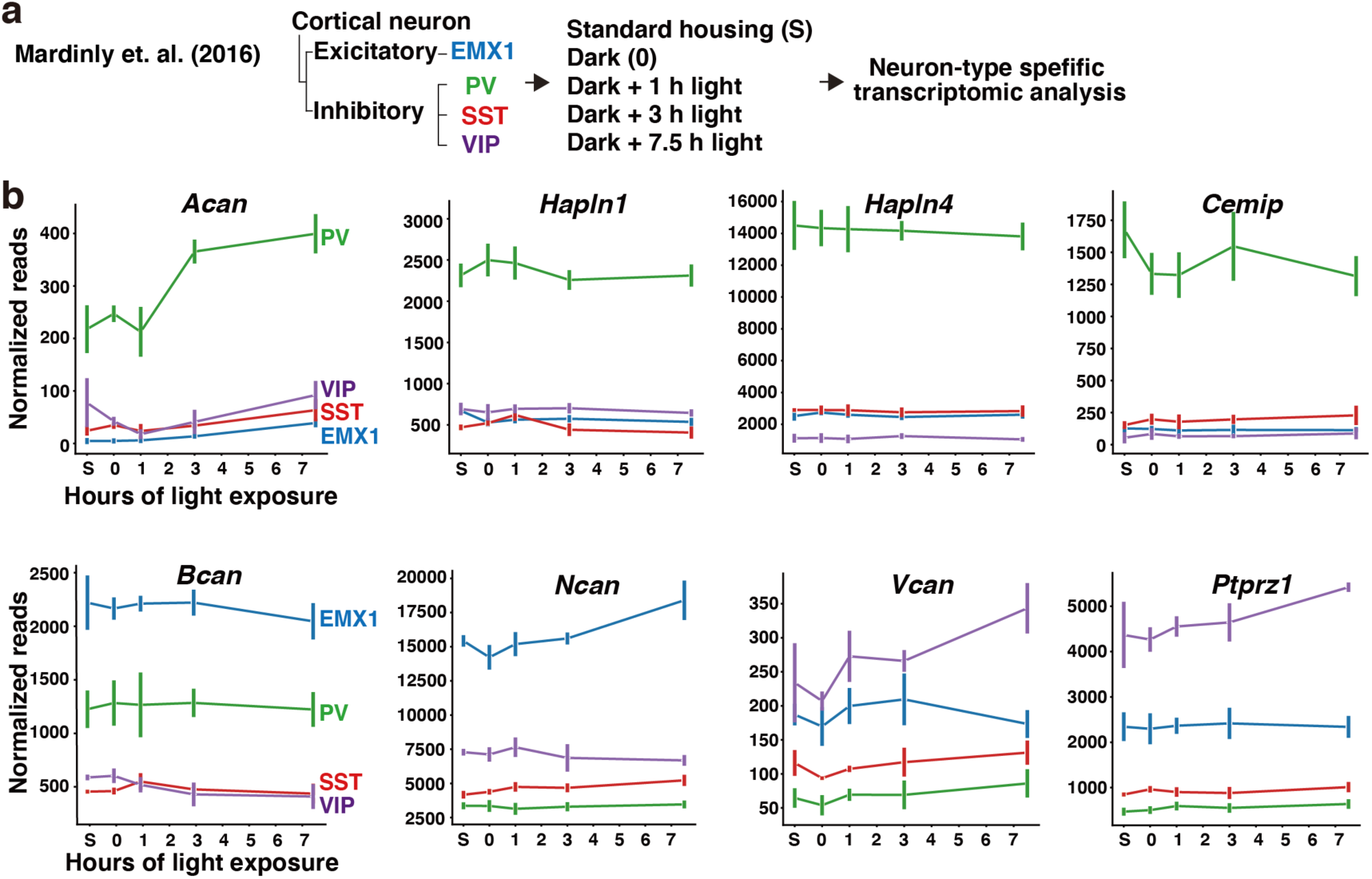
Activity-dependent *Acan* expression in PV neurons in the mouse visual cortex. (**a**) Experimental design of Mardinly et al. (2016) ^39^. Mice were housed in complete darkness for two weeks and then exposed to light stimulation for 0 to 7.5 hours to activate neurons in the visual cortex. Gene expression analysis was subsequently performed in EMX1-positive excitatory neurons and PV-, VIP-, and SST-positive inhibitory neuron subtypes. (**b**) Expression levels of PNN-related genes in EMX1-positive excitatory neurons (blue) and PV- (green), VIP- (purple), and SST-positive (red) inhibitory neuron subtypes following light exposure (n = 3 mice). Data are presented as mean ± SEM

### Bidirectional regulation of *Acan* expression by neuronal activity

Since increased neuronal activity enhances *Acan* expression, we next tested whether reduced activity has the opposite effect. To this end, we chronically suppressed neuronal firing using tetrodotoxin (TTX), a sodium channel blocker, and assessed the expression of PNN-related genes. We found that *Acan* expression in TTX-treated neurons was reduced to approximately 10% of control levels (Fig. 3a). *Acan* was the only PNN-related gene that exhibited bidirectional regulation, being upregulated by depolarization and downregulated by activity silencing. We also observed a significant reduction of aggrecan at the protein level by silencing neuronal activity with TTX (Fig. 3b, c). Compared to IEGs such as *Fos* and *Npas4*, which declined rapidly within hours, *Acan* expression showed a delayed decline over 24 hours after TTX treatment (Fig. 3d). To determine whether the reduction in *Acan* expression is associated with impaired PNN formation, we stained cortical neurons with WFA and observed a marked decrease in WFA intensity (Fig. 3e f). Taken together, these results indicate that *Acan* expression and subsequent PNN formation are bidirectionally regulated by neuronal activity.

**Figure 3.**
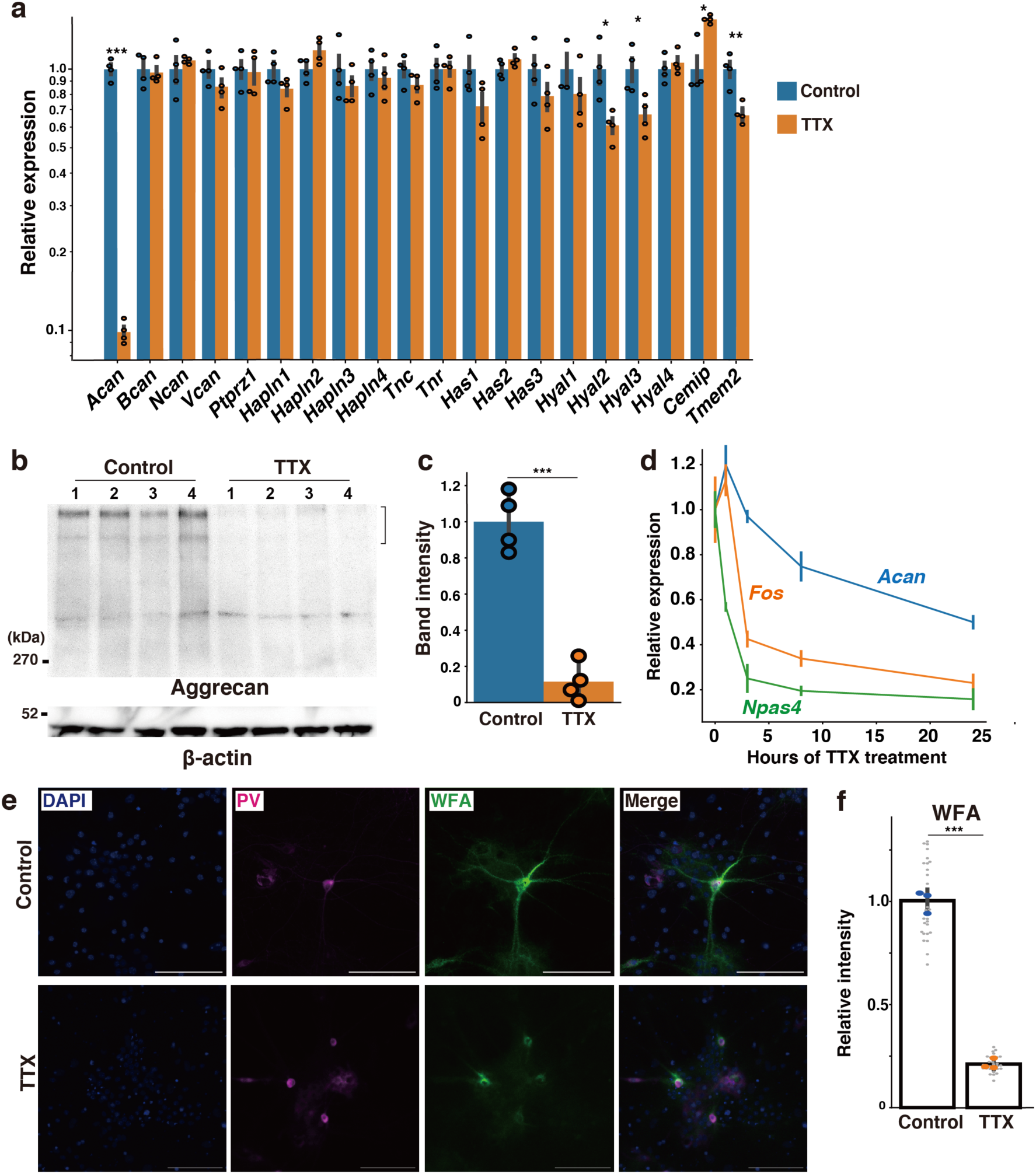
Bidirectional regulation of *Acan* expression by neuronal activity. (**a**) Expression levels of PNN-related genes in control and TTX-treated (2 µM, 48 h) cultures at DIV17, as determined by quantitative PCR. Values are expressed relative to control (n = 4 wells). (**b**) Western blot analysis of aggrecan and β-actin in control and TTX-treated (2 µM, 5 days) cultures at DIV14. (**c**) Band intensity of aggrecan shown in (b), normalized to β-actin and expressed relative to control (n = 4 wells). (**d**) Expression changes of *Acan*, *Fos*, and *Npas4* at the indicated time points after TTX treatment (n = 4 wells). (**e**) Representative images of WFA-positive PNNs surrounding PV neurons in control and TTX-treated (2 µM, 5 days) cultures at DIV14. (**f**) Relative WFA intensity around PV neurons. Gray dots represent individual images from each replicate, and blue (control) and orange (TTX) dots indicate the mean WFA intensity per well (n = 3 wells). All statistical tests are two-tailed Student’s t-tests. Data are presented as mean ± SEM. ***P < 0.001, **P < 0.01, *P < 0.05. Scale bars are 100 μm.

### CREB activation is essential for activity-dependent *Acan* expression and PNN formation

To elucidate the intracellular signaling mechanisms responsible for activity-dependent *Acan* expression, we pharmacologically dissected the upstream pathways in cultured cortical neurons. Inhibition of L-VGCCs with diltiazem markedly suppressed *Acan* mRNA expression (Fig. 4a), suggesting that calcium influx through these channels is essential for *Acan* transcription. In contrast, the expression levels of other CSPG genes remained unchanged after L-VGCC inhibition, underscoring the selective regulation of *Acan* by activity-dependent calcium signaling. Blocking NMDA and AMPA receptors with D-AP5 and NBQX also significantly reduced *Acan* expression (Fig. 4b), indicating that synaptic activity is required for its upregulation.

**Figure 4.**
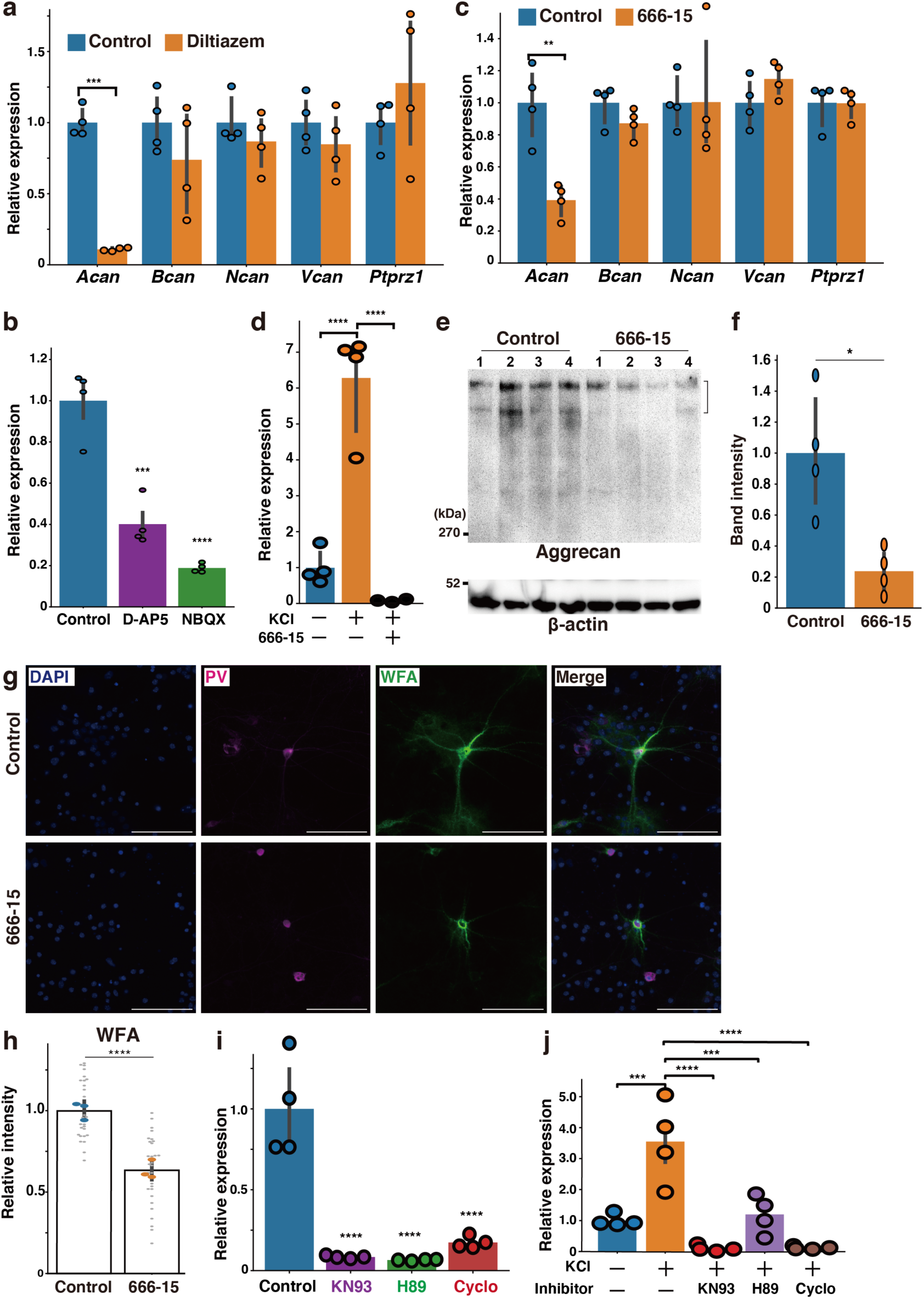
CREB activation is essential for activity-dependent *Acan* expression and PNN formation. (**a**) Expression levels of CSPG genes in control and diltiazem-treated (20 µM, 24 h) cultures at DIV18, as determined by quantitative PCR. Values are expressed relative to control (n = 4 wells). (**b**) Expression levels of *Acan* in control cultures and cultures treated with D-AP5 (20 µM, 24 h) or NBQX (20 µM, 24 h) at DIV17. Values are expressed relative to control (n = 4 wells). (**c**) Expression levels of CSPG genes in control and 666-15-treated (2 µM, 24 h) cultures at DIV16. Values are expressed relative to control (n = 4 wells). (**d**) Expression levels of *Acan* were analyzed at DIV16 in control and 666-15–treated (2 µM, 24 h) cultures followed by KCl-induced depolarization (25 mM, 24 h). Values are normalized to control (n = 4 wells). (**e**) Western blot analysis of aggrecan and β-actin in control and 666-15-treated (2 µM, 5 days) cultures at DIV14. (**f**) Band intensity of aggrecan shown in (e), normalized to β-actin and expressed relative to control (n = 4 wells). (**g**) Representative images of WFA-positive PNNs surrounding PV neurons in control and 666-15-treated (2 µM, 5 days) cultures at DIV14. (**h**) Relative WFA intensity around PV neurons. Gray dots represent individual images from each replicate, and blue (control) and orange (666-15) dots indicate the mean WFA intensity per well (n = 3 wells). Control cultures are the same as in Fig. 3f. (**i**) Expression levels of *Acan* in control cultures and cultures treated with KN-93, H89, or cyclosporin A (cyclo) (all 10 µM, 48 h) at DIV17. Values are expressed relative to control (n = 4 wells). (**j**) Expression levels of *Acan* were analyzed at DIV16 in control and KN-93, H89, or cyclo-treated (all 10 µM, 24 h) cultures followed by KCl-induced depolarization (25 mM, 24 h). Values are normalized to control (n = 4 wells). All statistical tests are two-tailed. Student’s t-tests were used in (a, c, e, f, h), and Dunnett’s tests in (b, d, i, j). Data are presented as mean ± SEM. ****P < 0.0001, ***P < 0.001, **P < 0.01, *P < 0.05. Scale bars are 100 μm.

**Figure 5.**
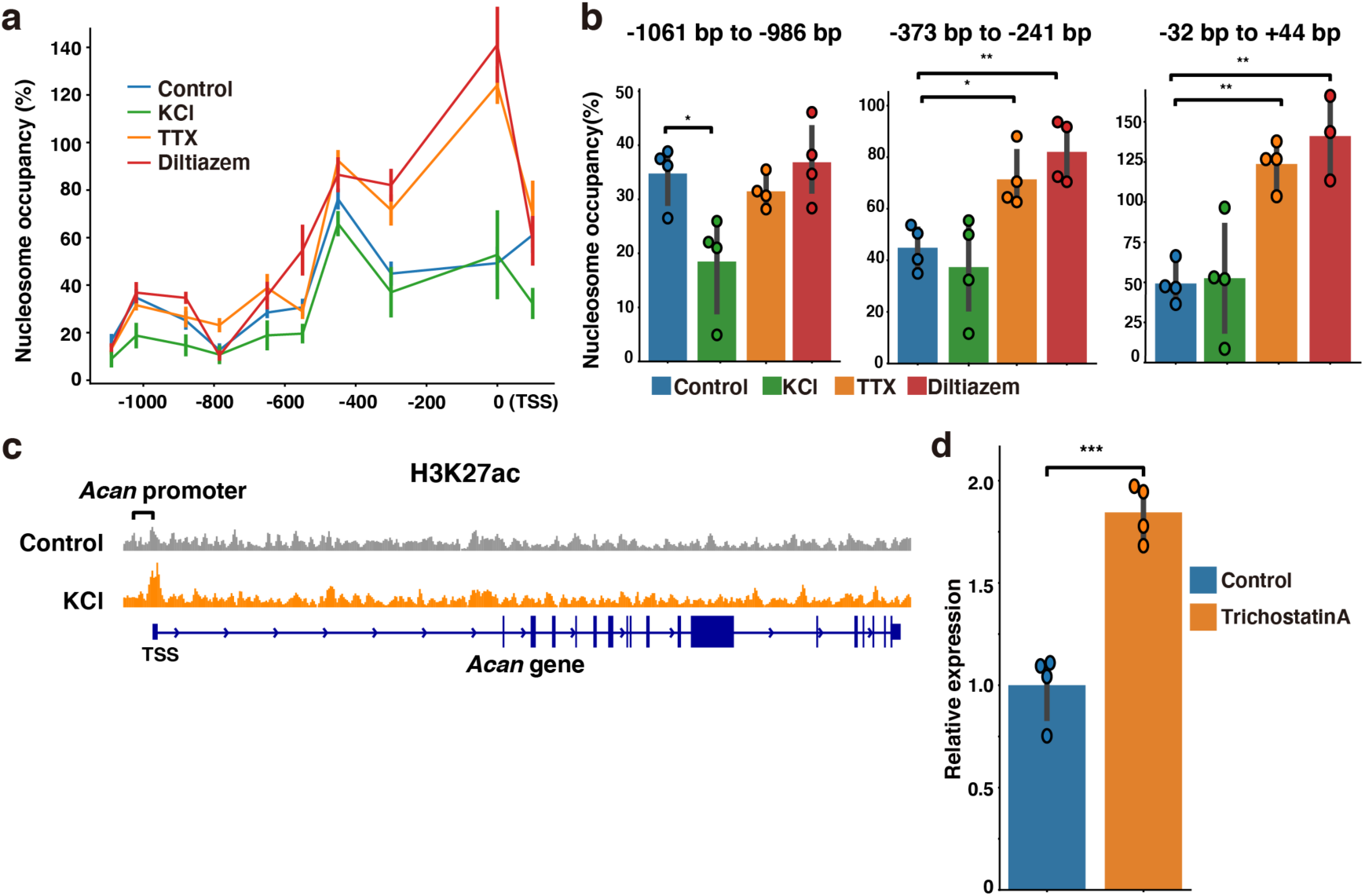
Neuronal activity induces chromatin remodeling at the *Acan* promoter. (**a**) Relative nucleosome occupancy upstream of the *Acan* TSS, covering up to 1000 bp, in control cultures (blue) and cultures treated with KCl (25 mM, 24 h; green), TTX (2 µM, 24 h; orange), or diltiazem (20 µM, 24 h; red). Values were normalized to those obtained from genomic DNA prepared without micrococcal nuclease digestion (n = 4 wells). (**b**) Relative nucleosome occupancy at three regions upstream of the *Acan* TSS (−1061 to −986 bp, −373 to −241 bp, and +1 to +44 bp) in control (blue) and KCl (green), TTX (orange), or diltiazem (red)-treated cultures (n = 4 wells). (**c**) IGV genome browser visualization of H3K27ac ChIP-seq signals at the *Acan* gene before (gray) and 2 h after KCl treatment (orange), based on reanalysis of published data from Malik et al. (2014) ^12^. (**d**) Expression levels of *Acan* in control and trichostatin A-treated (300 nM, 24 h) cultures at DIV17, as determined by quantitative PCR. Values are normalized to control (n = 4 wells). All statistical tests are two-tailed. Dunnett’s test was used in (b), and Student’s t-test in (d). Data are presented as mean ± SEM. ***P < 0.001, **P < 0.01, *P < 0.05.

We next examined whether activation of CREB, a key downstream effector of calcium signaling, is required for *Acan* expression. Treatment with the CREB inhibitor 666-15 significantly decreased *Acan* mRNA levels (Fig. 4c). The expression levels of other CSPG genes remained unchanged, indicating the specificity of CREB action. Furthermore, inhibition of CREB almost completely suppressed *Acan* induction by KCl-induced depolarization, indicating its essential role in activity-dependent transcription (Fig. 4d). The amount of aggrecan at the protein level was also significantly reduced after CREB inhibition. (Fig. 4e, f). To determine whether CREB-mediated *Acan* regulation is associated with PNN formation, we stained neurons with WFA and found that 666-15 treatment resulted in a 40% reduction in WFA intensity around PV neurons compared to controls (Fig. 4g, h).

CREB is a downstream target of several calcium-dependent signaling pathways, including CaMKs, PKA, and calcineurin. Inhibition of any of these pathways using KN-93 for CaMKs, H-89 for PKA, or cyclosporin A for calcineurin significantly reduced *Acan* mRNA levels under basal and depolarizing conditions, suggesting that multiple signaling cascades contribute to the maintenance of *Acan* expression (Fig. 4i, j). Taken together, these results indicate that activity-dependent *Acan* induction is mediated by calcium influx through L-VGCCs and subsequent activation of CREB, which in turn drives PNN formation around PV neurons.

### Neuronal activity induces chromatin remodeling at the *Acan* promoter

Neuronal activity regulates gene expression through epigenetic mechanisms, including chromatin remodeling, which modulates the accessibility of transcription factors. To investigate whether neuronal activity alters chromatin accessibility at the *Acan* promoter, we measured nucleosome occupancy using micrococcal nuclease digestion followed by quantitative PCR. Suppressing neuronal activity with TTX or inhibiting calcium influx with diltiazem increased nucleosome occupancy from the transcription start site (TSS) to approximately 300 bp upstream of the *Acan* promoter (Fig. 4a, b). In contrast, depolarization with KCl reduced nucleosome density approximately 1 kb upstream, indicating chromatin relaxation (Fig. 4a, b).

Next, we examined whether this activity-dependent chromatin remodeling is associated with histone acetylation, specifically H3K27 acetylation (H3K27ac), a hallmark of transcriptionally active chromatin. Reanalyzing previously published ChIP-seq data ^12^ revealed a significant increase in H3K27ac at the *Acan* promoter after depolarizing neurons with KCl (Fig. 4c). Consistent with this finding, treating cultured cortical neurons with the histone deacetylase (HDAC) inhibitor trichostatin A significantly increased *Acan* mRNA expression, indicating that activity-induced histone acetylation promotes *Acan* expression (Fig. 4d).

## Discussion

Our study reveals that *Acan* expression is induced in a neuronal activity-dependent manner in PV neurons, promoting PNN formation in the mouse cerebral cortex. We demonstrate that *Acan i*nduction requires sustained neuronal activity and is regulated by the CREB signaling cascade and chromatin remodeling. These findings unveil a novel molecular mechanism linking prolonged neuronal activity to the formation of PNNs, which stabilize neural circuits.

IEGs, such as *c-Fos* and *Npas4*, are rapidly induced by synaptic activity and do not require *de novo* protein synthesis ^7^. Many IEGs encode transcription factors that regulate LRGs, which are characterized by slower induction kinetics and dependence on newly synthesized proteins. Unlike IEGs, which are transiently activated, LRG induction requires sustained neuronal activity ^10^. Consistent with this finding, our results demonstrate that *Acan* expression requires prolonged stimulation over several hours rather than brief depolarization. This delayed transcriptional response likely reflects a functional role. First, neuronal activity promotes synaptic plasticity, which reorganizes neural circuits. After a delay, PNN formation occurs to stabilize the newly established connections. Thus, activity-dependent PNN formation may consolidate the reorganized circuit while inhibiting further plasticity.

We reveal that *Acan* is selectively induced in PV neurons in the mouse cerebral cortex. This selective expression may contribute to the formation of PNNs around these neurons. IEGs are widely expressed in various types of neurons, whereas LRGs are more specific to particular subtypes of neurons ^39,40^. For instance, *Npas4* is induced by neuronal depolarization in both excitatory and inhibitory neurons but activates distinct LRG programs in each cell type ^41^. One possible source of these differences is subtype-specific CREB signaling. PV neurons differ from excitatory neurons in that they use γCaMKI rather than γCaMKII for nuclear signaling ^42^. Additionally, PV neurons have low levels of CaMKIV, which contributes to their slow, sigmoidal CREB phosphorylation kinetics. These specialized signaling dynamics likely enable PV neurons to selectively induce specific LRGs, such as *Acan*, contributing to precise, subtype-specific transcriptional responses.

Additionally, recent transcriptomic and chromatin profiling studies have revealed substantial variations in the epigenomic landscapes of different neuronal subtypes, which contribute to their unique, activity-dependent transcription ^43–45^. One key mechanism involves histone acetylation, particularly H3K27ac, which facilitates the recruitment of RNA polymerase II to promoters and enhancers, thereby enabling transcriptional activation ^8^. We found that neuronal activity increases H3K27ac at the *Acan* promoter region. Consistent with this finding, treatment with the HDAC inhibitor significantly increased *Acan* expression in cultured cortical neurons. This finding supports a model in which activity-driven chromatin remodeling promotes *Acan* transcription and subsequent PNN formation. A recent study reported that deleting *Hdac2* selectively in PV neurons reduces *Acan* expression and impairs PNN formation, thereby attenuating fear memory recovery ^46^. However, given the functional diversity among HDAC isoforms, the effects of broad-spectrum inhibition by trichostatin A may differ from those of *Hdac2*-specific deletion.

In the developing brain, neurocan and tenascin-C bind to HA to form an embryonic-type ECM structure that supports neuronal proliferation, differentiation, and migration ^47–49^. After birth, the expression of neurocan and tenascin-C declines, whereas the expression of aggrecan, brevican, tenascin-R, and HAPLNs increases, promoting a transition from embryonic to adult-type ECM such as PNNs ^18,50^. Of the genes examined, only *Acan* exhibited bidirectional, activity-dependent regulation. Previous studies have shown that the expression of *Bcan* and *Hapln1* is modulated by neuronal activity ^36,38^. However, their changes appear to be more modest compared to the robust regulation observed for *Acan*. We previously reported that sulfation patterns of CSPGs play a crucial role in PNN formation and neuronal plasticity ^51,52^. Further research is needed to determine if neuronal activity modulates the sulfation structure of CSPGs in addition to the aggrecan core protein.

While the present study focused on the mechanisms underlying activity-dependent *Acan* gene expression, recent studies have established the critical role of *Acan* in PNN formation and the regulation of neuronal plasticity. Deletion of *Acan* in PV neurons disrupts PNN integrity, resulting in the persistence of critical period-like plasticity and improved cognitive performance ^53^. Similarly, adult-specific, virus-mediated knockout of *Acan* enhances visual cortical plasticity ^54^. In addition, deletion of *Acan* has also been reported to impair contextual fear memory ^55^, suggesting that the effects of *Acan* deletion may vary depending on the affected brain region and the type of learning task.

Alterations in PNNs have been implicated in various neurodevelopmental and neurodegenerative disorders. Methyl-CpG-binding protein 2 (MeCP2), a key epigenetic regulator mutated in Rett syndrome, influences PNN development. *Mecp2*-deficient mice exhibit precocious PNN formation and accelerated maturation of PV neurons ^56^. In schizophrenia, a significant reduction in aggrecan-positive PNNs has been observed in the amygdala ^57^, indicating compromised structural stabilization of neural circuits. Disruption of PNNs has also been reported in Alzheimer’s disease models, where abnormally activated microglia degrade PNN components including aggrecan ^58^. These findings suggest that PNN formation is involved not only in normal brain maturation but also in various brain disorders. Further research is needed to understand how the dysregulation of activity-dependent *Acan* expression contributes to disease pathophysiology.

## Acknowledgments

This research was funded by the Ministry of Education, Culture, Sports, Science & Technology, Japan [grant number 18K06130, 21H05681, and 24K09377 to S.M.], the Sumitomo Foundation [grant name Grant for Basic Science Research Projects to S.M.], and the LOTTE Foundation [grant name Lotte Shigemitsu Prize to S.M.]. The authors thank Seiya Kawano for technical advice on Western blotting, and the members of the Smart-Core-Facility Promotion Organization at Tokyo University of Agriculture and Technology for their assistance with confocal microscopy.

## Author contributions

K. N., A.M., and S.M. designed research, conducted experiments, and interpreted data. K. N. and S.M. wrote the draft, and all authors revised and approved the final manuscript.

## Materials and Methods

### Mice used in the present study

All animal experimental studies were conducted with the approval of the Committee on Animal Experiments of Tokyo University of Agriculture and Technology (Approval Code: R02-111, R03-103, R04-220). Pregnant ICR (RRID: MGI:5462094) were purchased from Japan SLC (Shizuoka, Japan).

### Primary cultures of cortical neurons

Cortical neuron cultures were prepared from E17 mouse embryos. After dissecting the brains from the fetuses, the meninges were carefully removed, and the cerebral cortices were dissected under a binocular stereomicroscope. All subsequent procedures were performed in a laminar flow hood. The tissue was cut into small pieces with scissors and washed three times with Hank’s Balanced Salt Solution (HBSS, Gibco) containing 10 mM HEPES (Gibco). Papain (Worthington Biochemical) was dissolved in 5 mL of HBSS and gently mixed by inversion. The solution was pre-incubated at 37 °C for 10 minutes. The tissue was then digested with 20 U/mL papain at 37 °C for 10 minutes. The enzymatic reaction was stopped by adding 500 µL of culture medium consisting of Neurobasal Plus Medium (Gibco) supplemented with 2% B-27™ Plus Supplement (Gibco), 5% horse serum (Sigma), and 1% penicillin/streptomycin (Sigma). The tissues were further dissociated by adding an appropriate amount of DNase and triturating them with a Pasteur pipette. The cell suspension was filtered through a cell strainer and centrifuged at 400×g for 4 minutes at room temperature. After discarding the supernatant, 2 mL of fresh culture medium was added and the number of cells was counted. The cells were plated at a density of 2 × 10⁵ cells per well onto glass coverslips in 4-well plates. The coverslips were pre-coated with a 500 µL solution of 0.1 mg/mL poly-L-ornithine (Wako) per well. One-third of the culture medium was replaced with serum-free culture medium every three days. To prevent glial overgrowth, the cultures were treated with 3 µM cytosine β-D-arabinofuranoside at DIV4.

### Pharmacology

Neurons were treated at DIV9-18. To induce neuronal activity, the cultures were stimulated with 25 mM KCl for 1, 3, 8, or 24 hours. To silence neuronal activity, the neurons were treated with 2 μM TTX (Nacalai Tesque) for 1, 3, 8, 24, or 48 hours, or for 5 days. Additional cultures were treated with 20 μM D-AP5 (R&D Systems), 20 μM NBQX (Cayman Chemicals), or 20 μM diltiazem (Wako Chemicals) for 24 or 48 hours. To inhibit CREB-mediated transcription, neurons were treated with 2 μM 666-15 (MedChemExpress) for 24 or 48 hours or 5 days. Other cultures were treated with 10 μM KN-93 (Selleck), 10 μM H89 (Selleck), or 10 μM cyclosporin A (LKT Laboratories) for 24 or 48 hours. To inhibit HDAC, neurons were treated with 300 nM trichostatin A (Wako) for 24 hours. All concentrations listed above refer to final concentrations. The specific duration of each treatment is indicated in the relevant figures and figure legends.

### Quantitative PCR

Total RNA was extracted from primary cultured cortical neurons using the RNeasy Mini Kit (Qiagen). Complementary DNA was synthesized using the PrimeScript RT Reagent Kit (Takara Bio). Real-time quantitative PCR was performed using SYBR Premix Ex Taq (TakaRa Bio) on a Thermal Cycler Dice Real Time System III (Takara Bio). All primer sequences are listed in the Key Resources Table. Relative gene expression levels were calculated using the comparative cycle threshold (Ct) method and normalized to β-actin (*Actb*) expression.

### Western blotting

Primary cultured cortical neurons were lysed in PBS containing 1% Triton X-100 and a protease inhibitor cocktail (Sigma), and incubated on ice for 30 minutes. For chondroitinase ABC digestion, the lysates were treated with 5 milliunits of Proteus vulgaris chondroitinase ABC (Sigma) at 37 °C for 2 hours. The digested lysates were then denatured with 2% sodium dodecyl sulfate and 5% β-mercaptoethanol at 60 °C for 20 minutes. The proteins were separated using either 4.5% or 10% polyacrylamide gel electrophoresis and transferred to polyvinylidene difluoride membranes. The membranes were blocked with 2% skim milk in PBS containing 0.1% Tween 20 (PBST) and incubated overnight at 4 °C with the following primary antibodies: anti-aggrecan (rabbit, 1:2,000; Sigma) and anti-β-actin (mouse, 1:10,000; Proteintech). After washing, the membranes were incubated with the appropriate horseradish peroxidase (HRP)-conjugated secondary antibodies (1:2000) for 1 hour at room temperature. Protein bands were visualized using the Immobilon Western Chemiluminescent HRP Substrate (Millipore) and imaged using a LuminoGraph I system (ATTO).

### Cell staining

Primary cultured cortical neurons were fixed with 4% paraformaldehyde in PBS for 15 minutes, permeabilized with 0.25% Triton X-100 in PBS, and blocked with PBST containing 2% bovine serum albumin (Sigma). The neurons were incubated overnight at 4 °C with biotinylated WFA (1:1000, Vector Laboratories) and an anti-parvalbumin antibody (mouse; 1:2000, Swant). After washing, the neurons were incubated with Alexa Fluor 647-conjugated anti-mouse IgG antibody and Alexa Fluor 488-conjugated streptavidin (Thermo Fisher), followed by nuclear staining with DAPI (0.1 μg/mL). For each condition, three to four wells were prepared and stained. Ten images were acquired per well using an AX R confocal microscope using a 20× objective lens. (Nikon). Imaging was performed

Image analysis was performed using ImageJ to quantify WFA intensity on the soma of PV neurons. First, the PV signals were binarized to detect PV-positive somata. Using size thresholding, only soma-sized objects were selected, and binary masks of individual PV neuronal somata were generated. These masks were then overlaid onto the corresponding WFA-stained images, allowing measurement of WFA fluorescence intensity specifically on the PV neurons. Statistical analyses were performed using the mean WFA intensity per well as a single data point. Each condition included three to four biological replicates (wells).

### Nucleosome occupancy in the upstream region of the *Acan* Gene

Nucleosomes were isolated from primary cultured cortical neurons using the EpiScope Nucleosome Preparation Kit (Takara Bio). Briefly, primary cultured cortical neurons were lysed to isolate nuclei, which were then digested with micrococcal nuclease to selectively degrade linker DNA regions not protected by nucleosomes. After removing histones with proteinase K treatment, nucleosome-protected DNA fragments were purified. Quantitative PCR was performed using the purified nucleosomal DNA as a template to assess nucleosome occupancy in the upstream region of the *Acan* gene. All primer sequences are listed in the Key Resources Table. Relative nucleosome occupancy was calculated using the comparative Ct method. Ct values were normalized to those obtained from genomic DNA prepared without micrococcal nuclease digestion, which served as a reference for total DNA input.

### Reanalysis of *in vivo* RNA-seq and *in vitro* ChIP-seq data

The RNA-seq data from Mardinly et al. (2016) ^39^ were downloaded as Excel files from the Gene Expression Omnibus (GEO; accession number GSE77243). The data were then visualized as line plots using Python within the Jupyter Notebook environment. The ChIP-seq data Malik et al. (2014) ^12^ were downloaded as bigWig files from GEO (GSE60192) and visualized using the Integrative Genomics Viewer IGV with alignment to the mm9 genome build. All datasets were normalized to the total number of uniquely mapped reads.

### Quantification and statistical analysis

All data are presented as the mean ± standard error of the mean (SEM). Statistical analyses were performed using Python within the Jupyter Notebook environment and Easy R ^59^. For comparisons between two groups, unpaired, two-tailed Student’s t-tests were used. For comparisons involving multiple experimental groups against a single control group, a Dunnett’s test was performed. A p-value of less than 0.05 was considered statistically significant. Details of the statistical analyses, including the specific tests used, sample sizes (n), and P-values, are provided in the corresponding figures and figure legends.

---

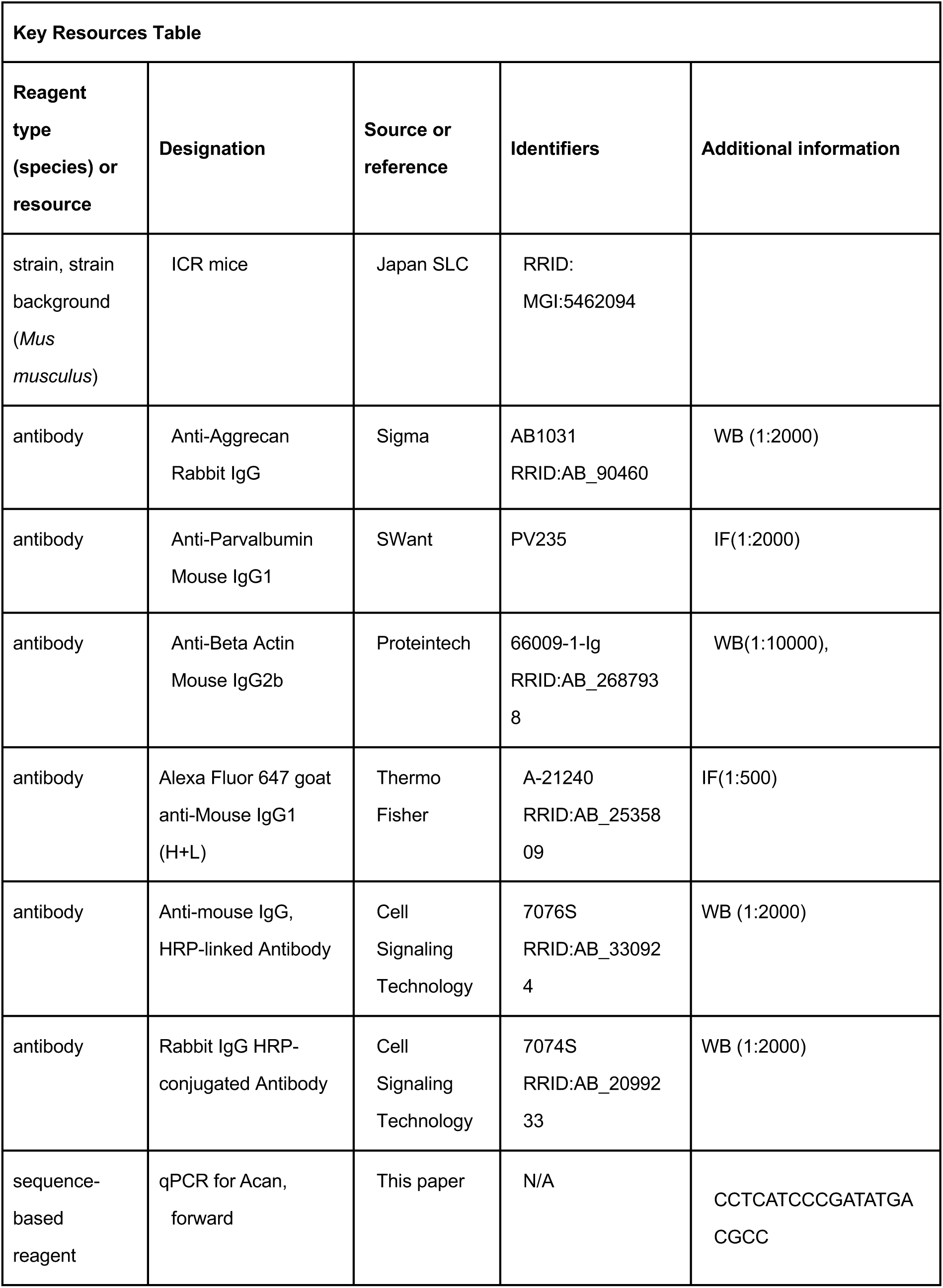
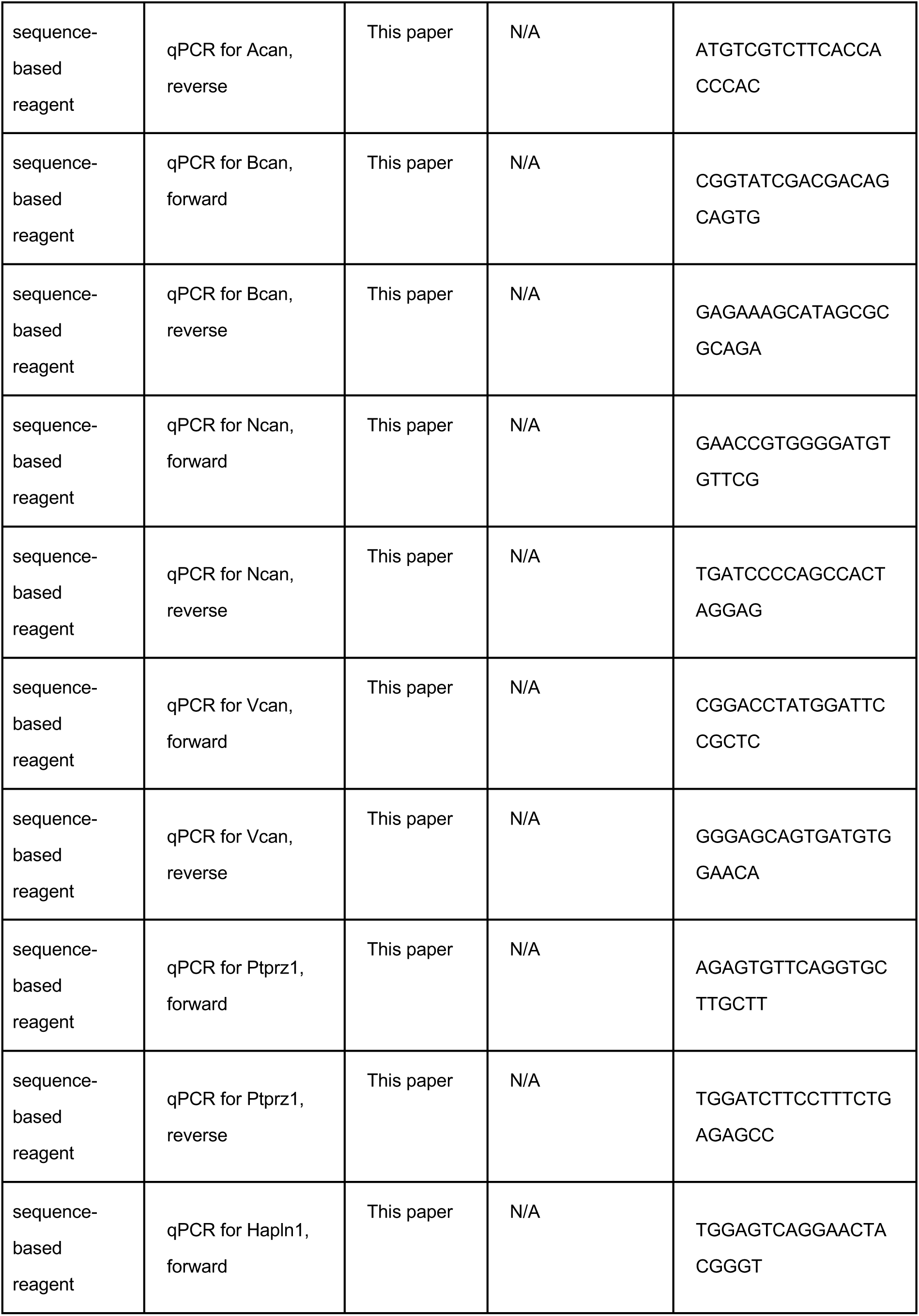
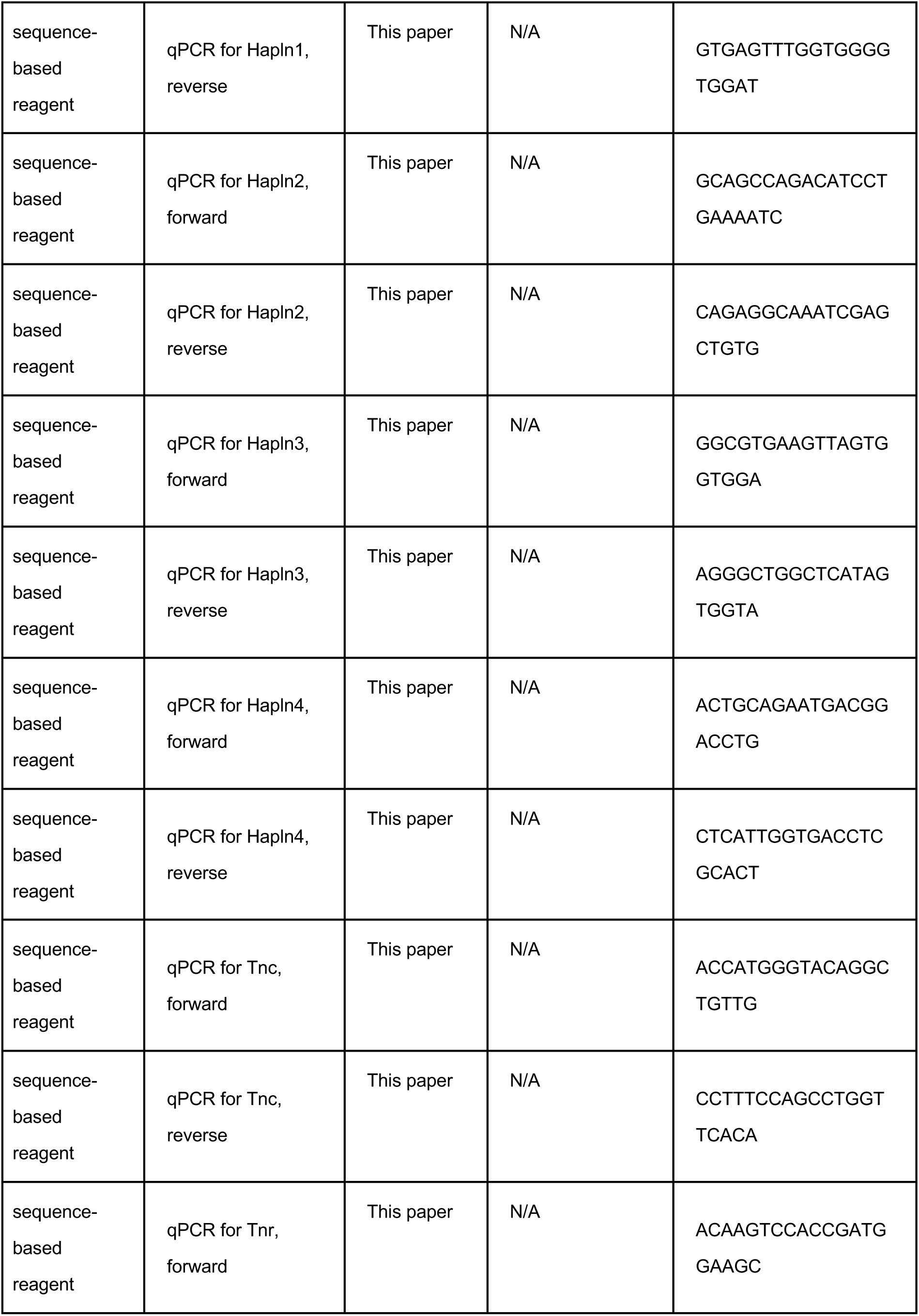
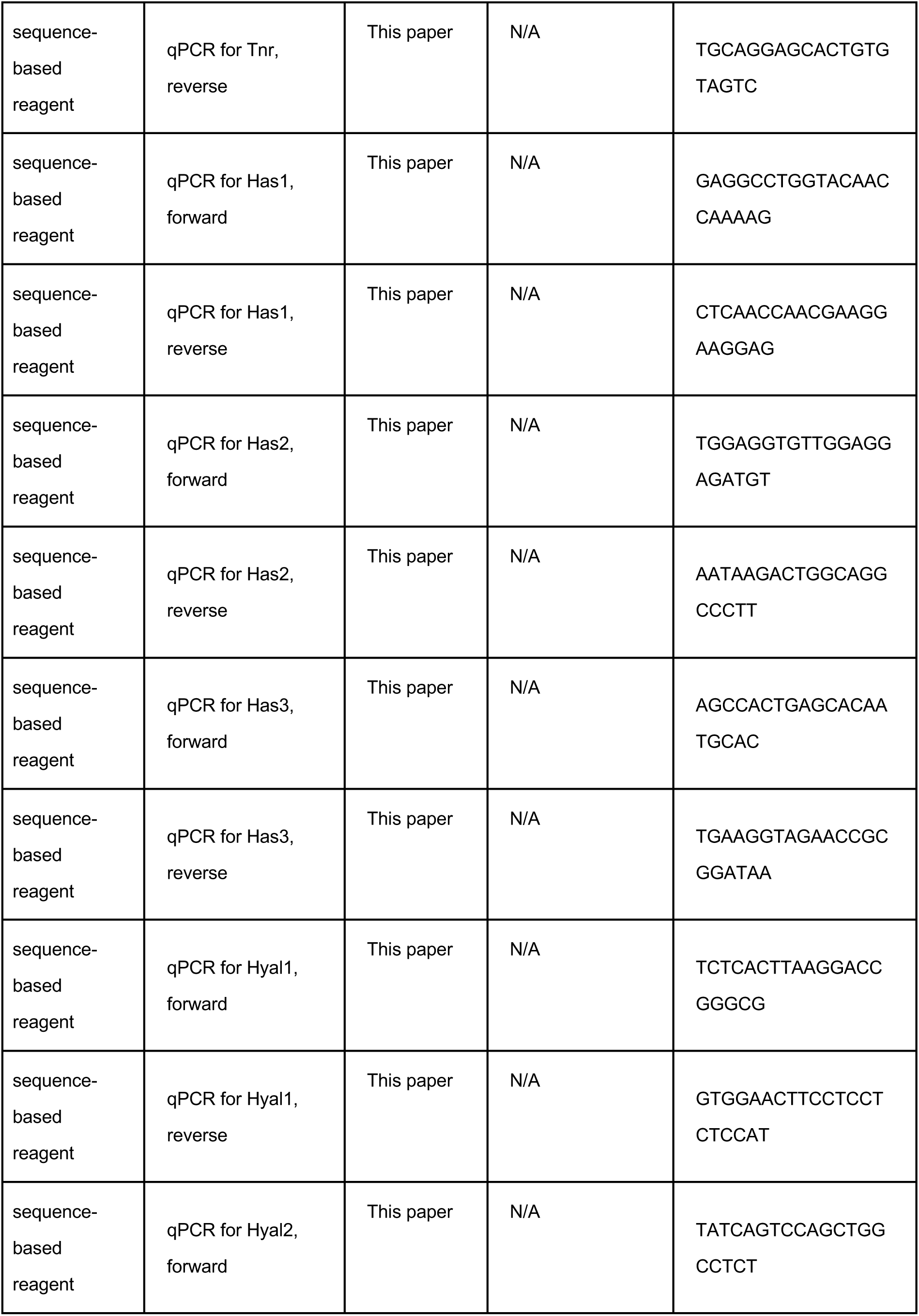
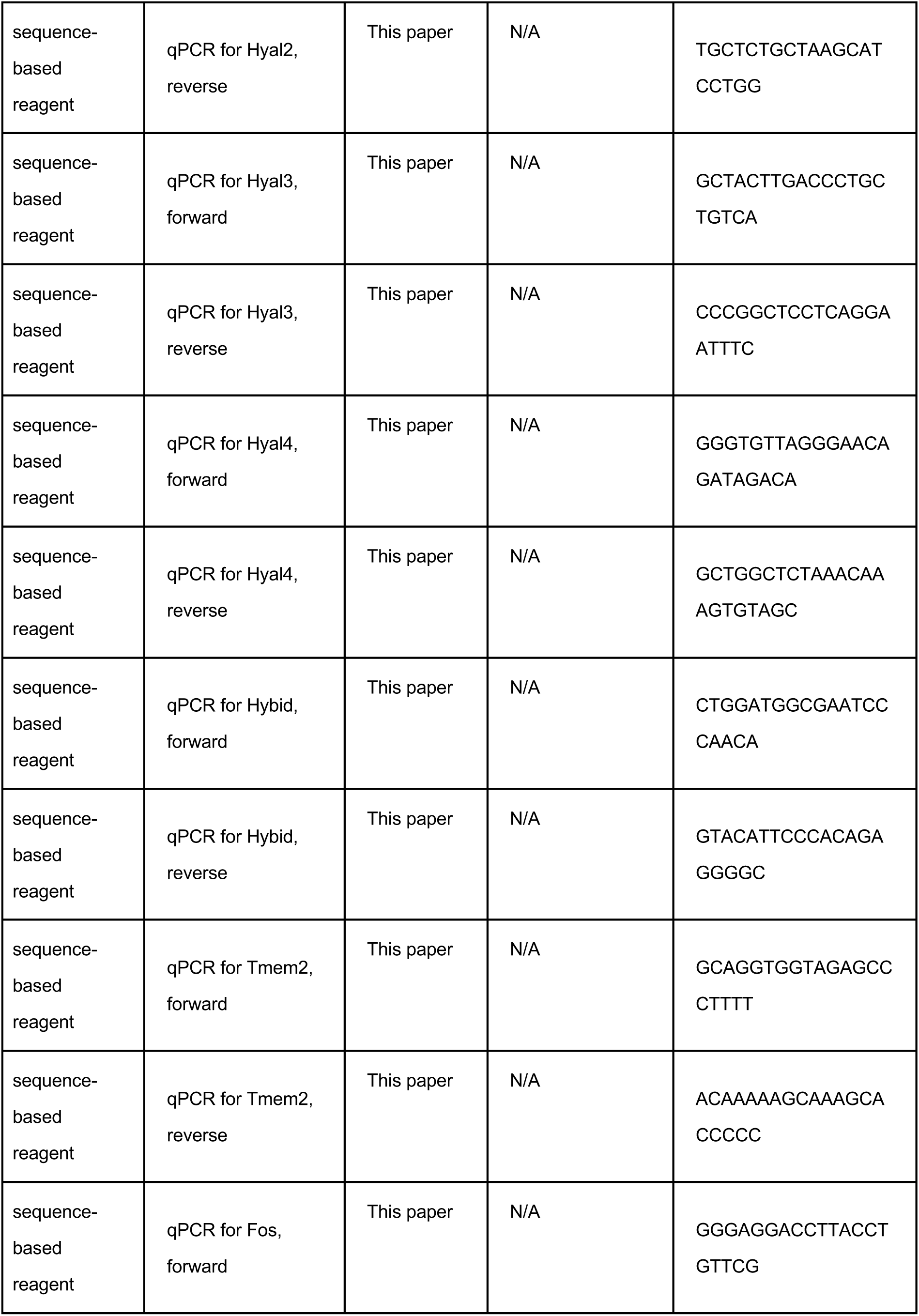
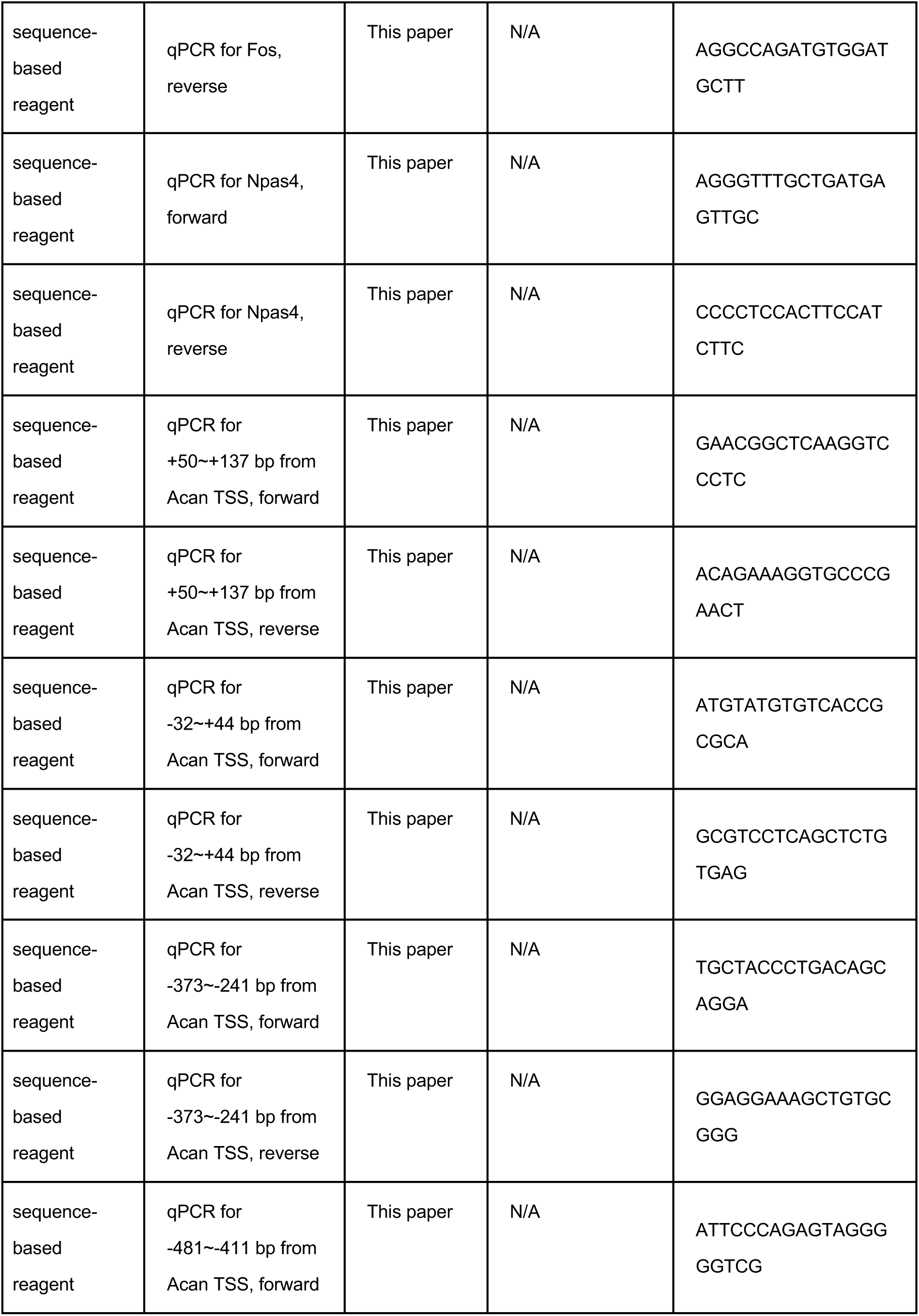
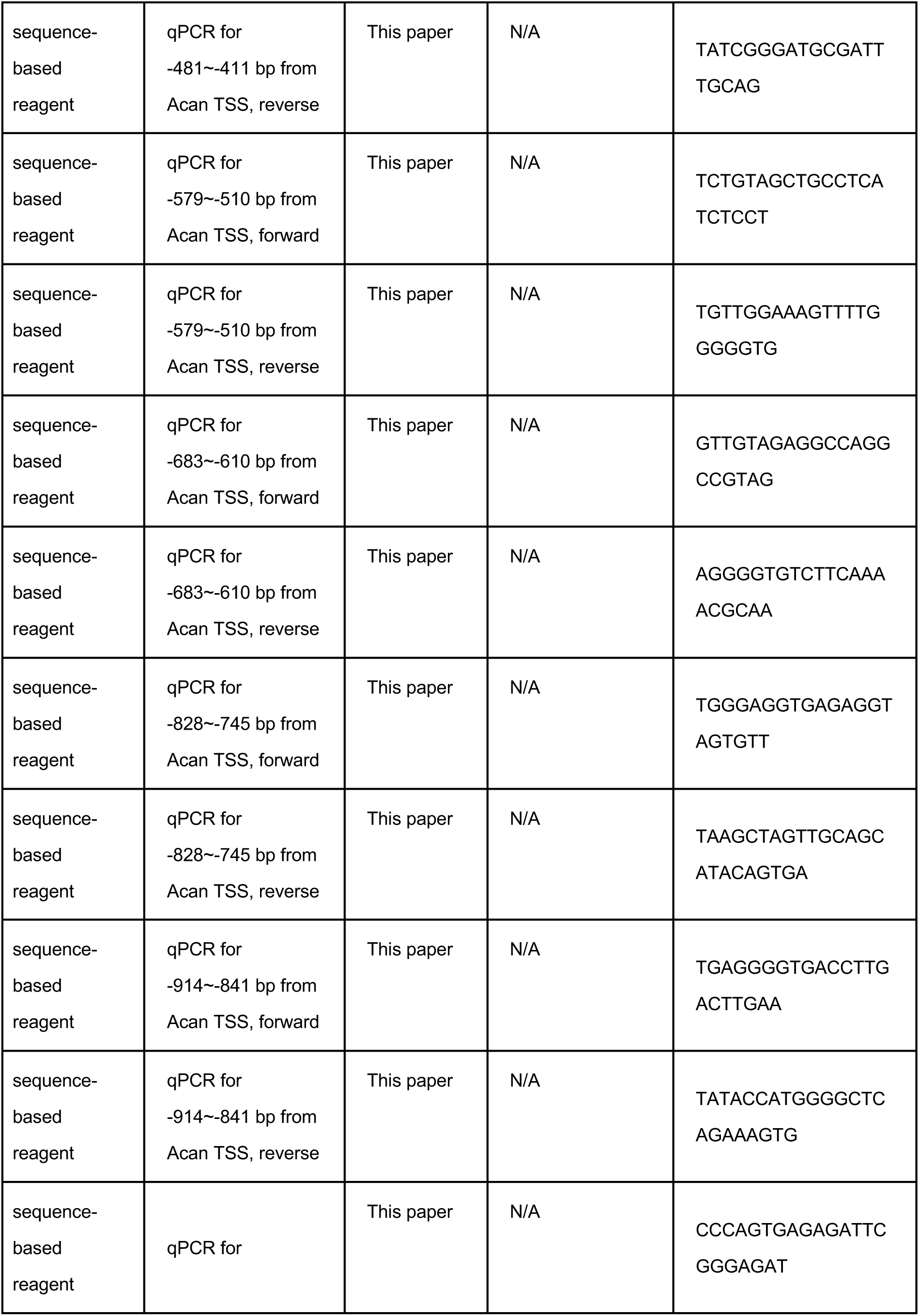
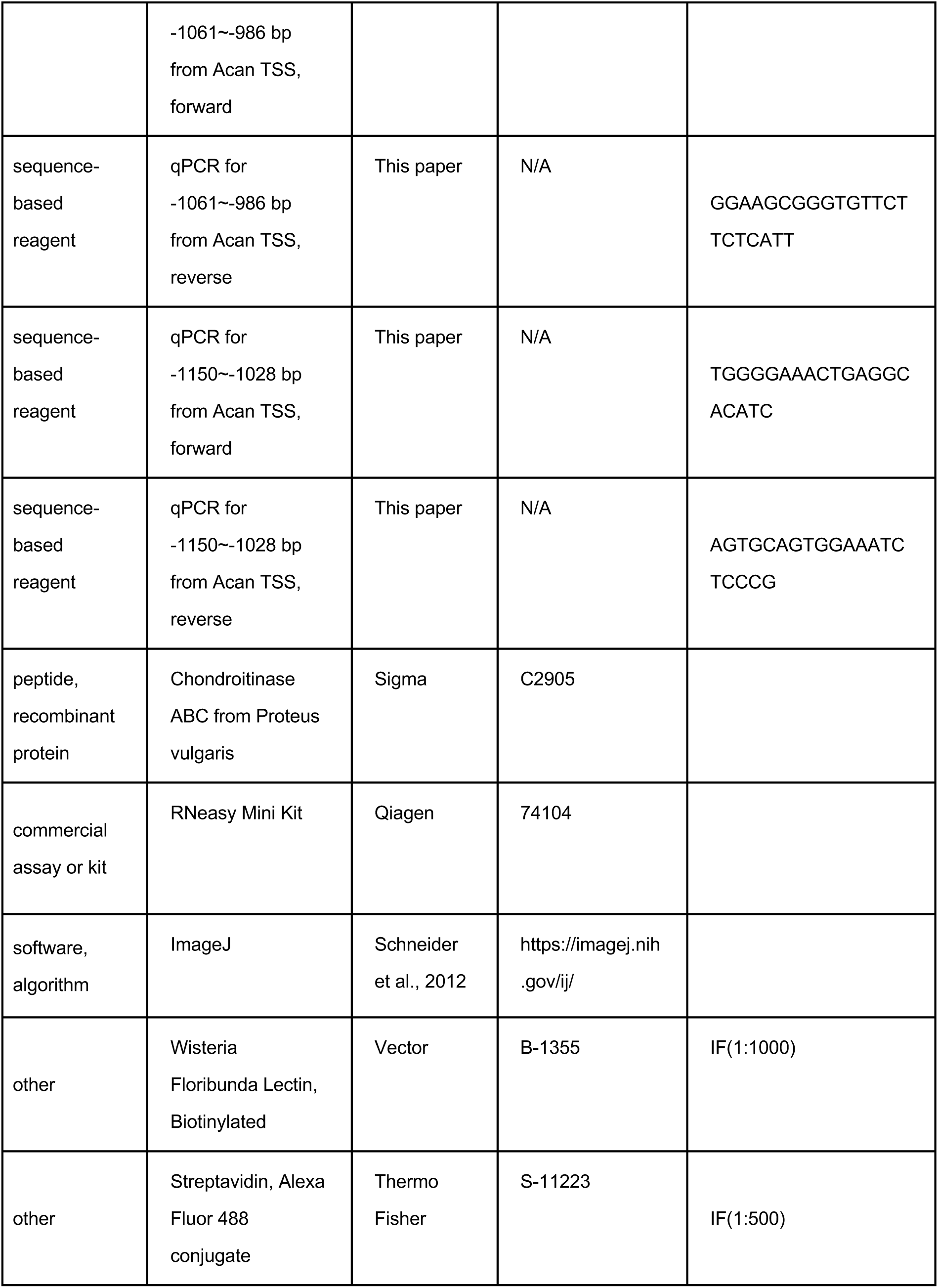
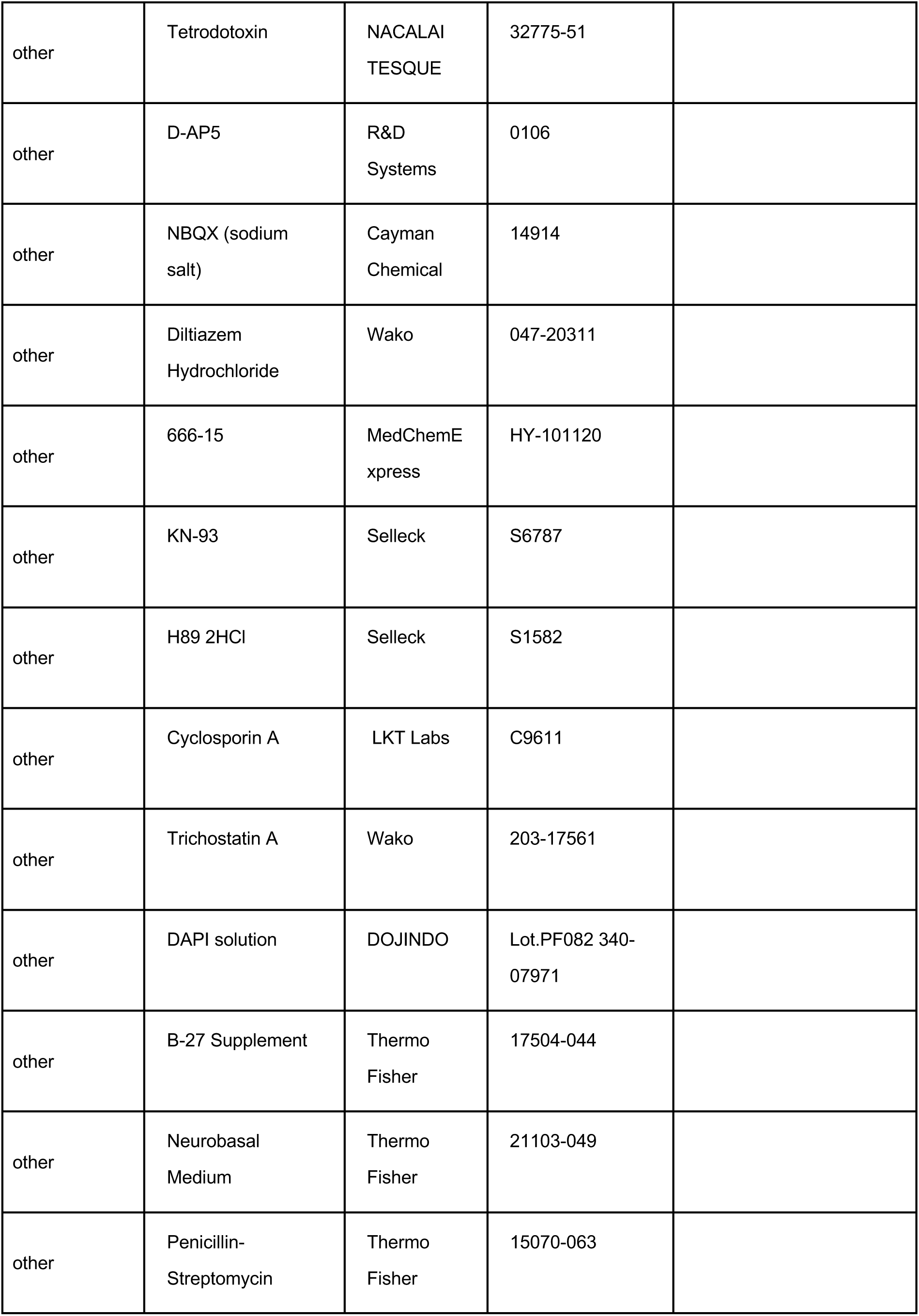
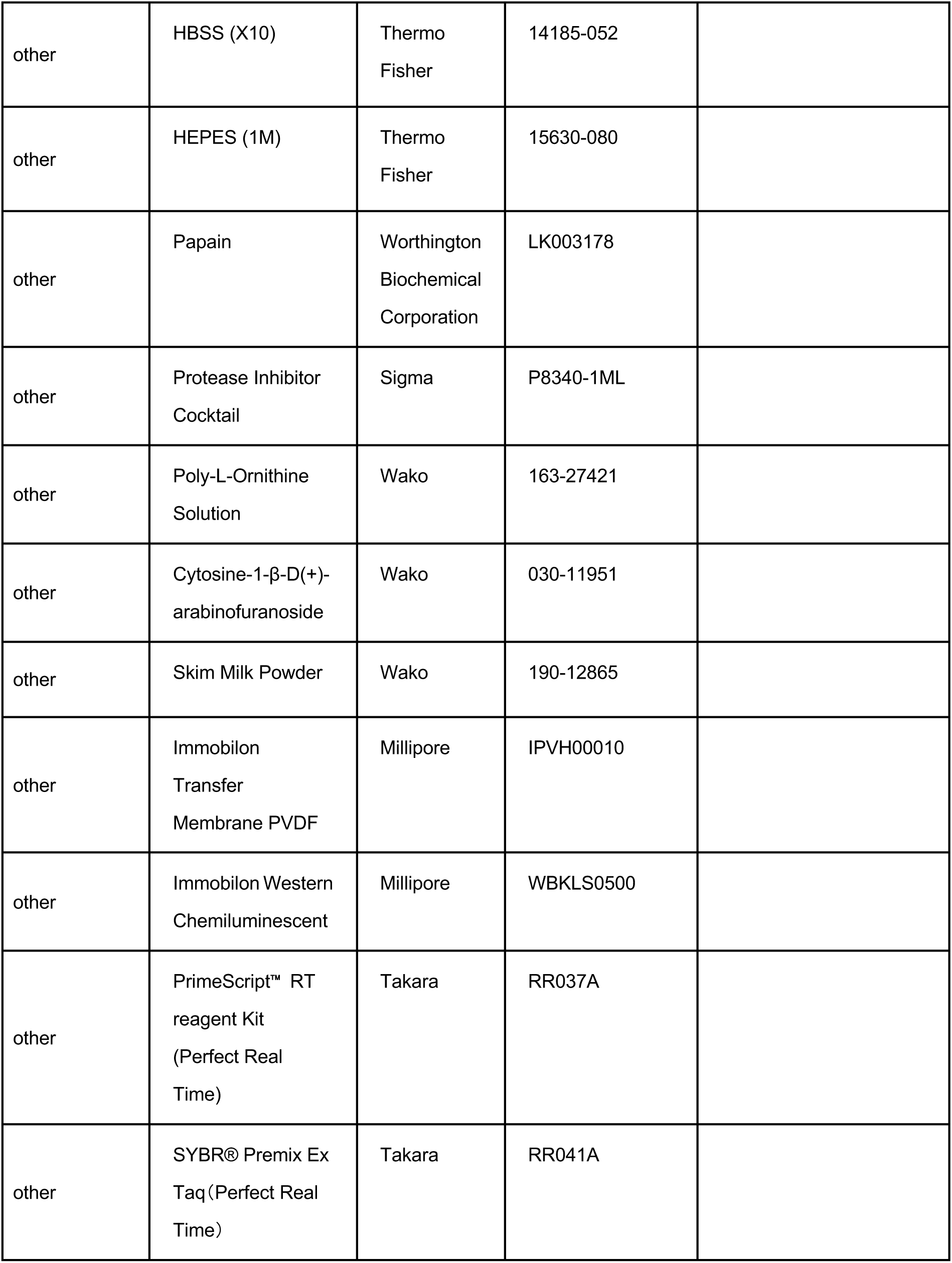
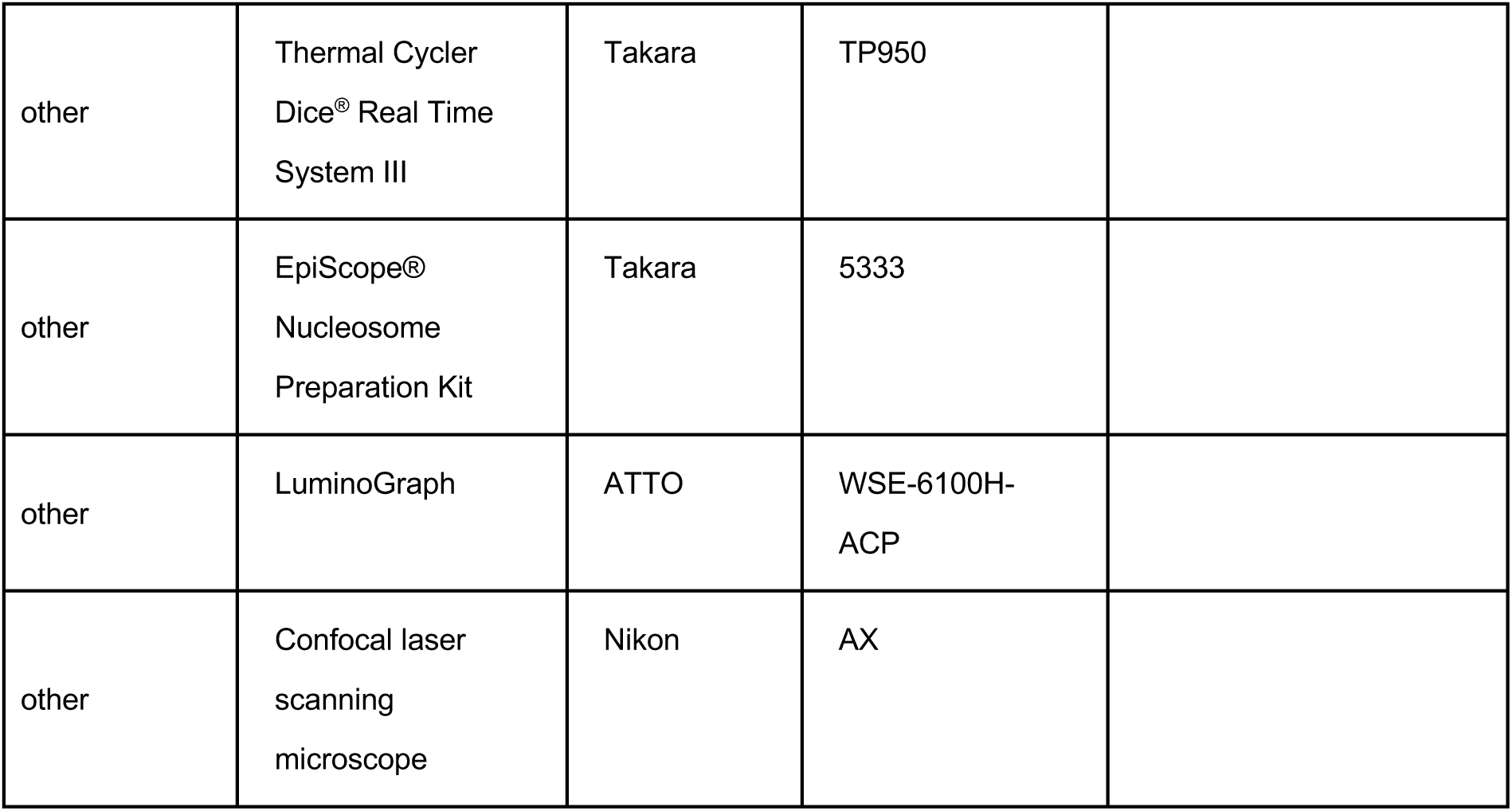

