## Supplementary data 1 for "Neuronal activity induces aggrecan expression to drive perineuronal net formation in cortical neurons"

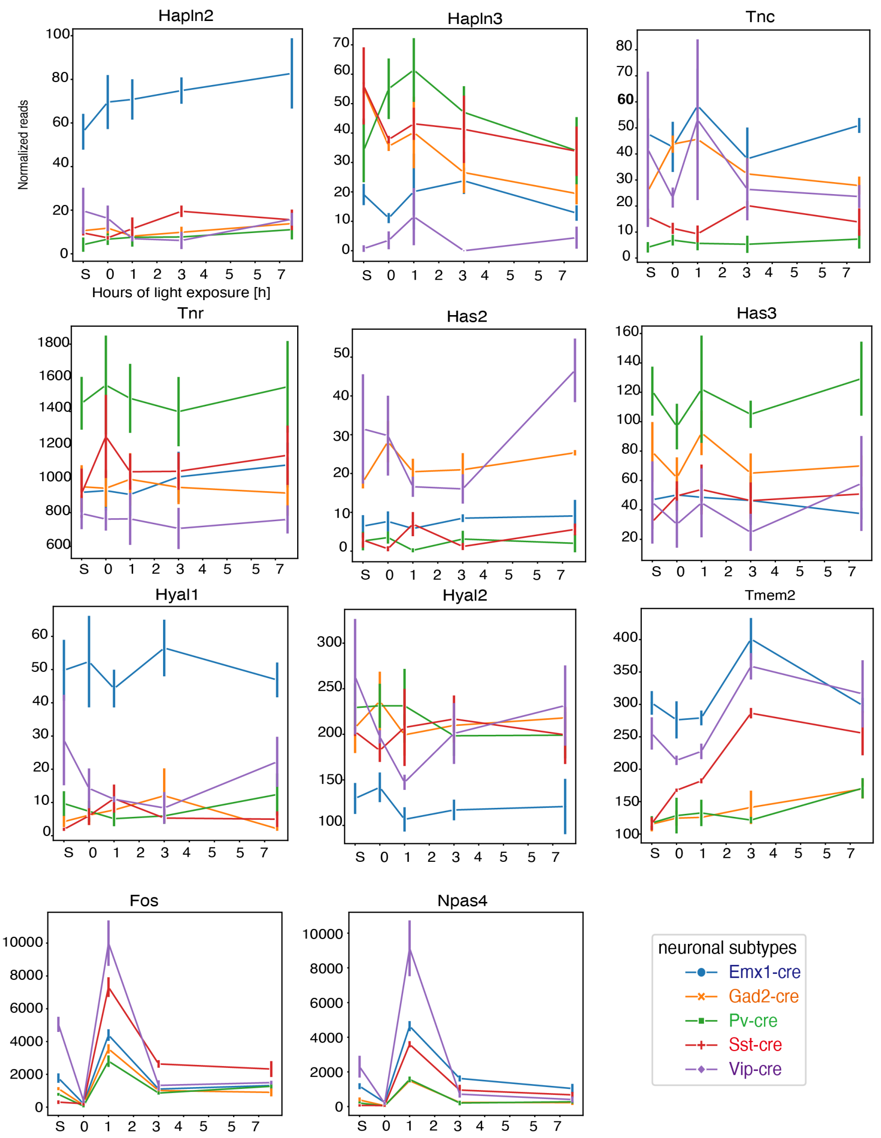


**Supplementary data 1.** Expression levels of PNN-related genes in EMX1-positive excitatory neurons (blue) and Gad2- (orange), PV- (green), VIP- (purple), and SST-positive (red) inhibitory neuron subtypes following light exposure (n = 3 mice). Data are presented as mean ± SEM
